# Cross-linked RNA Secondary Structure Analysis using Network Techniques

**DOI:** 10.1101/668491

**Authors:** Irena Fischer-Hwang, Zhipeng Lu, James Zou, Tsachy Weissman

## Abstract

Next generation sequencing and biochemical cross-linking methods have been combined into powerful tools to probe RNA secondary structure. One such method, known as PARIS, has been used to produce near base-pair maps of long-range and alternative RNA structures in living cells. However, the procedure for generating these maps typically relies on laborious manual analysis. We developed an automated method for producing RNA secondary structure maps using network analysis techniques. We produced an analysis pipeline, dubbed cross-linked RNA secondary structure analysis using network techniques (CRSSANT), which automates the grouping of gapped RNA sequencing reads produced using the PARIS assay, and tests the validity of secondary structures implied by the groups. We validated the clusters and secondary structures produced by CRSSANT using manually-produced grouping maps and known secondary structures. We implemented CRSSANT in Python using the network analysis package NetworkX and RNA folding software package ViennaRNA. CRSSANT is fast and efficient, and is available as Python source code at https://github.com/ihwang/CRSSANT.

## Background

The base-pairing properties of RNA enable its ability to form complex macro-molecular structures, which are critical to the variety of functions that different RNA perform as messengers, gene regulators, and subcomponents of complex molecular machinery and cellular networks [6, 10]. So crucial is structure to RNA function that a variety of techniques have been developed to probe the trajectory of an RNA molecule from single-stranded transcript to complex, folded macromolecule—plus all variety of stable and unstable intermediates and alternative conformations that might be adopted in between. Predicting the base-pairing of RNA nucleotides, or RNA secondary structure, has long been the goal of algorithms that calculate possible free-energy structures [26] or exhaustively search for shared structural motifs in nucleotide sequences [11]. Meanwhile, insights into the three-dimensional, or tertiary structure of RNA, have predominantly been obtained using x-ray crystallography and nuclear magnetic resonance spectroscopy [4].

In the last two decades, a host of chemical methods have dramatically advanced both secondary structure prediction and tertiary structure probing [33]. Despite these advancements, the structural insights are often limited, for example, to providing information about which bases are paired but without specifics about their pairing partners, or to capturing ensemble effects averaged over a large number of molecules [21]. However, recent work based on in vivo crosslinking and next-generation sequencing provides a new method for identifying base-pairing interactions in living cells [3, 30, 23, 17, 13, 31, 27, 36]. One such method, psoralen analysis of RNA interactions and structures—PARIS for short—targets base-paired regions of RNA and uses sequencing to directly read out paired bases. First, a psoralen derivative is used to cross-link base pairs in RNA helices in living cells. Next, RNase digestion is used to remove regions of RNA that are single-stranded. Cross-linked base pairs are then ligated, denatured and reverse transcribed to produce a single DNA molecule that is complementary to the RNA stem comprising the original cross-linked RNA base pairs. Each DNA read is sequenced and mapped to a reference sequence, producing “gapped” reads, which are paired—duplexed—reads without single-stranded loop regions. Each gapped read contains a left and right portion—or “arm”—where “left” denotes the 5′, or upstream position in the reference genome and “right” denotes the 3′, or downstream position.

The gapped reads produced by the PARIS method make it possible to identify not only which bases in a sequence are involved in base-pairing, but also the counter-parties in each base pair. The insights into base-pairing provided by the PARIS method were used to validate structures obtained by other structure probing methods, and were also used to identify the structural basis for long-range RNA interactions in long non-coding RNA. Although the PARIS method is extremely powerful in revealing basic stem structures and long-range interactions, the process of clustering gapped reads in order to extrapolate underlying RNA structures is laborious. In this manuscript, we present a computational method dubbed CRSSANT, which is intended to be an analytical complement to the PARIS assay. CRSSANT leverages network analysis techniques—also frequently referred to as “graph” techniques—in order to automate analysis of sequencing reads produced by the PARIS assay.

In theory, it is straightforward to infer RNA base pairs from PARIS reads. Each gapped read was produced by subjecting a single RNA stem to a succession of biochemical processes that makes it possible to read out the nucleotides and base pairs underlying the stem. However, in practice a number of issues make it difficult to easily interpret PARIS reads. First, the cross-linker employed in the PARIS assay has a preference for staggered uridine bases, and, like any catalyst, is less than 100% efficient. Furthermore, certain RNA structures may even block cross-linking. In addition to issues with cross-linking, the ligation step is known to have a low efficiency of approximately 5%, at most. These complications mean that the PARIS method is unable to capture all RNA structures, making it difficult to directly infer properties like the relative abundance of alternative structures or long-range interactions [22]. Additionally, the RNase digestion process is known to be highly variable, which may sometimes result in reads with unexpectedly long arms.

To account for these issues, the CRSSANT pipeline comprises two main analysis procedures: clustering and structure extrapolation. It accepts as input gapped reads produced by the PARIS method that have been mapped to a reference sequence by the STAR read aligner [8]. First, the pipeline transforms reads into a network representation. This representation facilitates clustering of highly similar reads into duplex groups (DGs). DGs are further processed to produce stem groups (SGs), which are used to identify candidate regions of a reference sequence that could undergo base-pairing and give rise to an RNA stem. To produce SGs from DGs, reads with unusually long arms are removed from each DG. The remaining filtered reads are used to identify consensus regions of the reference sequence, which are tested for base-pairing ability using state-of-the art RNA structure prediction software. Finally, the pipeline outputs structure maps for predicted RNA stems as lists of predicted base pairs.

An overview of the PARIS assay and CRSSANT methodology is shown in Figure 1.

**Fig. 1.**
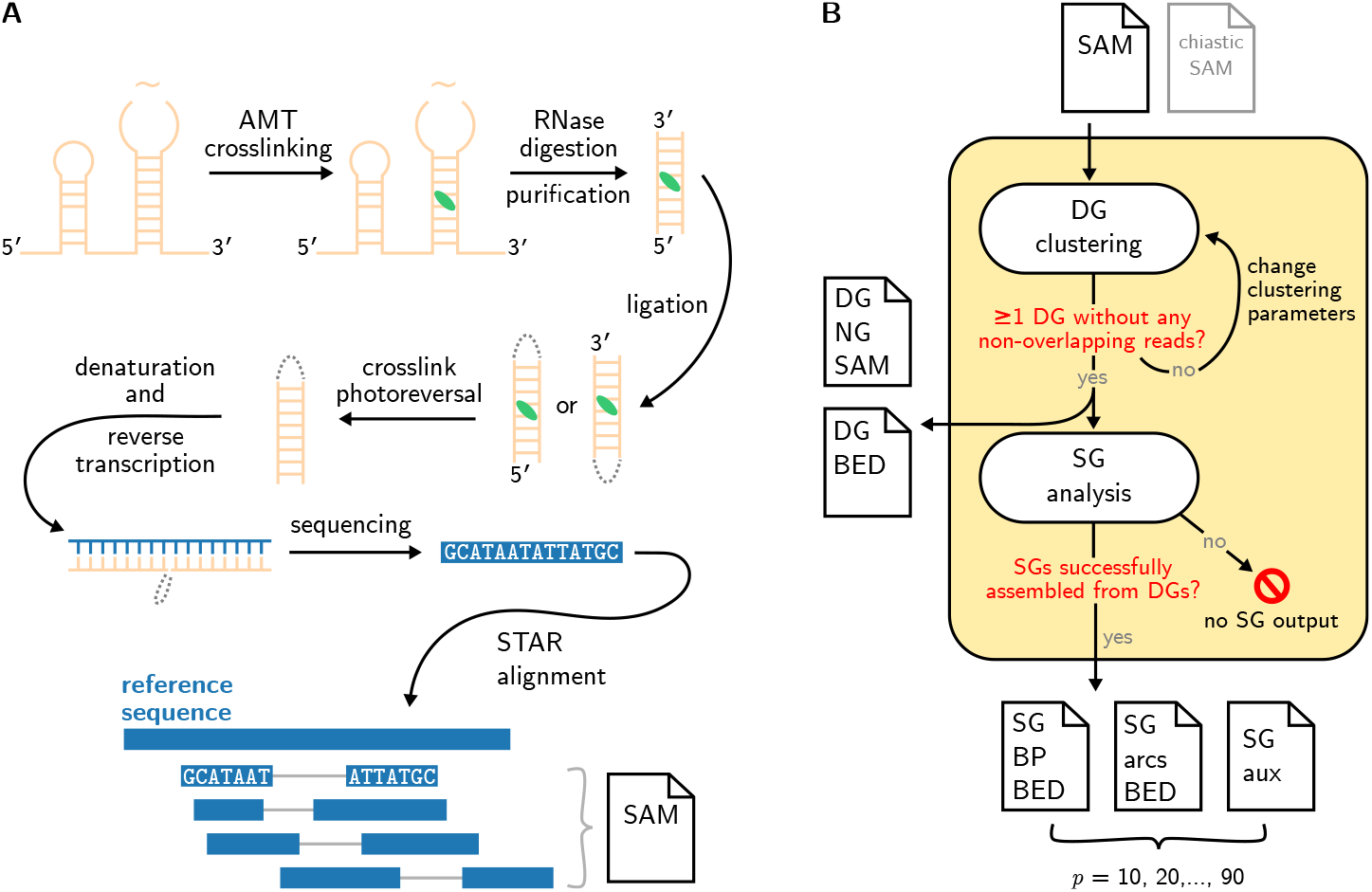
Overviews of CRSSANT and PARIS. (A) Cartoon overview of the PARIS assay with critical steps: in vivo cross-linking of RNA (beige) via psoralen-derivative 4′-aminomethyltrioxsalen (AMT, green lozenge), RNase digestion and purification, two possible ligation modes (dashed gray line), cross-link photoreversal to remove AMT cross-links, reverse transcription producing complementary DNA (blue), sequencing, and alignment to a reference sequence. Solid gray lines indicate paired read arms. (B) Cartoon overview of the CRSSANT analysis pipeline, which accepts as input a SAM file of aligned reads (and, if applicable, a SAM file of chiastic reads, gray) and performs the following steps: DG clustering (producing two DG files if successful), SG clustering (which follows from DG clustering if DG clustering is successful), and output SG files which are produced if SG analysis is successful. Output SG files are produced in sets of three, one for each SG assembly percentile threshold *p*.

**Fig. 2.**
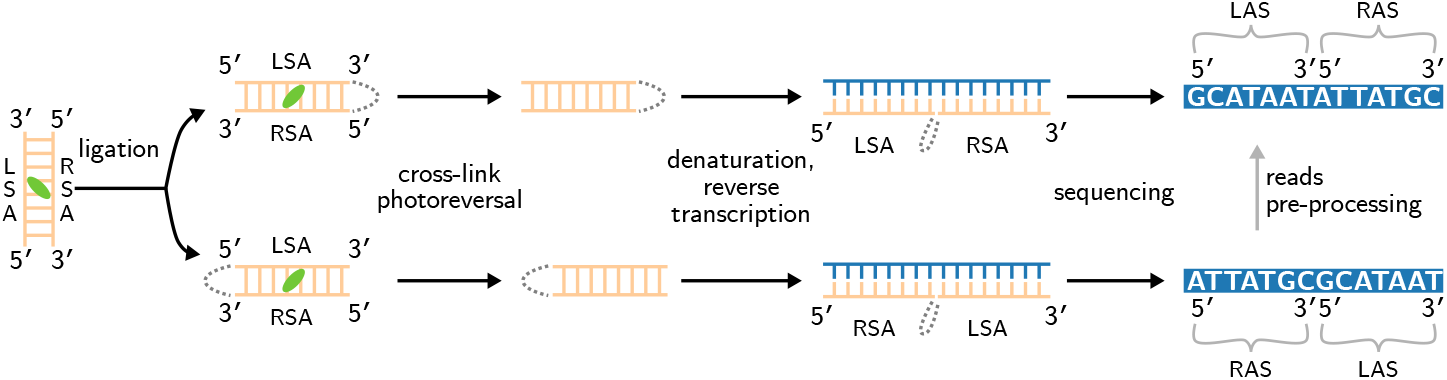
Ligation may lead to normal or chiastic gapped reads. A single cross-linked RNA stem (beige) comprising a left stem arm (LSA) and right stem arm (RSA) may undergo ligation at either of the stem ends. Ligation between the 3′ end of the left stem arm and the 5′ end of the right stem arm ultimately lead to a “normal” gapped read (top path). Reverse transcription of the ligated RNA stem arms produces a complementary DNA molecule (blue), which is sequenced to produce a read (blue bar). In a normal gapped read, the read comprises a sequence of nucleotides corresponding to the left read arm—or left arm sequence (LAS)—followed by a sequence of nucleotides corresponding to the right read arm—or right arm sequence (RAS). The entire read sequence (white letters in blue bar) is ordered such that the left arm sequence is 5′ of the right arm sequence (stem arm directions are indicated by the 5′, 3′ pairs). Ligation may also occur between the 5′ end of the left stem arm and the 3′ end of the right stem arm, resulting to a “chiastic” gapped read (bottom path). In a chiastic read, the left arm sequence is ordered 3′ of the reversed right arm sequence. The chiastic read sequence must be pre-processed in order to undergo the rest of the CRSSANT pipeline.

## Results

### DG assignment validation and choice of default clustering parameters

We tested DG clustering methods and set default clustering parameters for CRSSANT by validating CRSSANT DG assignments on sets of PARIS reads generated in [24]. These sets comprised reads that were mapped to the following human gene sequences: RNA component of mitochondrial RNA-processing endoribonuclease (RMRP), ribonuclease P RNA component H1 (RPPH1), subunits 18S, 5.8S and 28S from ribosomal ribonucleic acid (rRNA) transcription unit 45S, and small nuclear ribonucleic acids (snRNAs) U2, U4, U5, U6, U4atac, and U6ata. Basic properties of the datasets are summarized in Table 1.

**Table 1.** Datasets used in all experiments. Basic properties of the datasets used to evaluate CRSSANT, including haploid gene length (in bases), number of reads (Reads), average read length (also in bases), and average sequencing coverage [18].

| Dataset | Gene | Gene length | Reads | Average read length | Average sequencing coverage |
| --- | --- | --- | --- | --- | --- |
| RMRP | RMRP | 268 | 694 | 53 | 136 |
| RPPH1 | RPPH1 | 333 | 1,142 | 48 | 166 |
| rRNA | 18S | 1,870 | 5,670 | 73 | 220 |
|  | 5.8S | 158 | 252 | 69 | 109 |
|  | 28S | 5,071 | 3,975 | 66 | 52 |
| snRNA | U2 | 187 | 459 | 44 | 107 |
|  | U4 | 145 | 1,899 | 44 | 582 |
|  | U6 | 106 | 187 | 47 | 84 |
|  | U5 | 116 | 1,100 | 41 | 392 |
|  | U4atac | 130 | 165 | 47 | 60 |
|  | U6atac | 125 | 99 | 51 | 40 |
| snoRNA | U3 | 214 | 4,481 | 47 | 988 |

We tested two types of DG clustering methods: cliques-finding and spectral clustering. For both clustering methods we tested the effect of different overlap thresholds *t*_o_. We also tested two spectral clustering-specific parameters: eigenvalue test size *n*_eig_, and eigenratio threshold *t*_eig_. In brief, *n*_eig_ is the number of eigenvalues that are considered during spectral clustering, and *t*_eig_ is the threshold that the ratio of gaps between consecutive eigenvalues must exceed to merit a new cluster. For more details about these parameters, please refer to the Methods section. We tested various ranges of clustering parameters: for both cliques-finding and spectral clustering methods we tested *t_o_* from 0.1 to 0.9 at increments of 0.1, and for spectral clustering we tested *n*_eig_ = {10, 20, 100} and *t*_eig_ = {3, 5, 10}.

We relied on a combination of three indicators to judge CRSSANT DG assignments against those generated in [24], which serve as ground truth DG sets. The first indicator, which we call “mismatch fraction,” measures the amount of variation between the ground truth DG set and the CRSSANT DG assignments. Mismatch fraction is calculated by comparing the ground truth DG assignments with the CRSSANT DG assignments for each pair of reads that were assigned to valid DGs using the CRSSANT pipeline. A penalty is accrued whenever there is a difference between the ground truth and CRSSANT DG assignments, i.e. if the two reads have the same DG assignment in the ground truth but are assigned to different DGs by CRSSANT, or vice versa. To obtain the mismatch fraction, the total penalty is normalized by 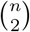, where *n* is the number of reads that were successfully assigned to valid DGs using the CRSSANT pipeline. In the best case, the mismatch fraction is zero.

The second indicator is the fraction of original reads that were not assigned to a valid DG using the CRSSANT pipeline. Ideally, this fraction is close or equal to zero, since all reads were assigned to DGs in the ground truth sets. Finally, the third indicator is the fraction of non-overlapping reads, or the fraction of reads that are assigned to the same DG that do not share any overlap in either arm. Because DGs are expected to contain highly similar reads, this number should be zero for the optimal clustering method. In fact, the cliques method of clustering DGs always results in zero non-overlapping reads, since every node in a clique must share an edge with every other node, i.e. each read corresponding to a node must overlap every other read more than *t*_o_.

The optimal clustering method and corresponding optimal parameter(s) were chosen to be those for which the fraction of non-overlapping reads equals zero, and the sum of the mismatch fraction and the fraction of unassigned reads—a quantity we refer to as the clustering score—is minimized. Read set statistics, optimal clustering method and parameters, mismatch fraction and fraction of unassigned reads are reported for each dataset in Table 2. The full set of results for DG clustering experiments is recorded in Additional file 1.

**Table 2.** Duplex group clustering results using default CRSSANT parameters: spectral clustering under *t*_o_ = 0.5, *n*_eig_ = 10 and *t*_eig_ = 5. The number of reads and ground truth duplex groups (GT DGs) are listed for each dataset. The resultant fraction of unassigned reads (FUR), mismatch fraction (MF) and number of non-overlapping reads are also listed.

| Dataset | Gene | Reads | GT DGs | FUR | MF | Non-overlapping reads |
| --- | --- | --- | --- | --- | --- | --- |
| RMRP | RMRP | 694 | 32 | 0.06 | 0.24 | 171 |
| RPPH1 | RPPH1 | 1,142 | 30 | 0.03 | 0.08 | 336 |
| rRNA | 18S | 5,670 | 266 | 0.04 | 0.03 | 2,538 |
|  | 5.8S | 252 | 7 | 0.01 | 0.28 | 126 |
|  | 28S | 3,975 | 287 | 0.07 | 0.01 | 531 |
| snRNA | U2 | 459 | 13 | 0.01 | 0.06 | 0 |
|  | U4 | 1,899 | 7 | 0 | 0 | 3 |
|  | U6 | 187 | 3 | 0.01 | 0.41 | 5 |
|  | U5 | 1,100 | 1 | 0 | 0 | 0 |
|  | U4atac | 165 | 3 | 0 | 0.01 | 0 |
|  | U6atac | 99 | 5 | 0.02 | 0.07 | 17 |

For optimal clustering parameters (Additional file 1, bold rows), the clustering score ranges from 0 in the case of snRNA U5 to 0.36 for RMRP, out of a maximum clustering score of two (since the clustering score is the sum of two values with maximum value 1). The perfect clustering score achieved by the DG clusters in the snRNA U5 dataset is due to the fact that the ground truth DG set contained only a single DG, as did the optimal spectral clustering. The results for the U4atac dataset are only slightly larger than 0.

The results further show that optimal clustering method and corresponding parameters varied from dataset to dataset, and indeed, from gene to gene within the same dataset. Cliques-finding was the optimal clustering method chosen for validation sets RMRP, rRNAs 18S and 28S, and snRNAs U4 and U6atac. For all other validation sets, spectral clustering was chosen as the optimal clustering method, and the optimal parameters *t*_o_ and *t*_eig_ vary. None of these trends can be explained by basic dataset attributes, including number of reads, reference sequence length spanned by the reads, nor the approximate coverage of the validation sets. With six of the 11 validation sets having optimal clustering results using spectral clustering, we set spectral clustering to be the default CRSSANT clustering method. Due to consistently good spectral clustering performance when *n*_eig_ = 10, we set as default *n*_eig_ to 10. Based on the median of the optimal parameters for the six validation sets that underwent spectral clustering, we also set as default *t*_eig_ = 5 and *t*_o_ = 0.5. When the cliques-finding method is selected, the pipeline uses a default value of 0.1 for *t*_o_. The results for DG clustering using the default clustering method and parameters are shown in Table 2.

### SG structure validation and prediction

After finding the optimal clustering method and parameters for creating DGs, we tested the SGs assembled from DGs for their potential to base pair and form stable RNA stems. In addition to the RMRP, RPPH1, rRNA and spliceosomal datasets, we tested an additional set of PARIS reads comprising gapped reads that mapped to the reference sequence encoding small nucleolar RNA U3 (snoRNA U3, see Table 1). For each of these datasets, there exist sets of structures based on a variety of structure-probing experimental methods [16, 34, 28, 2, 25], which were recorded as individual base pairs (Additional file 5). We used these known base pair sets to both validate known RNA structures as well as reveal possible alternative base-pairings.

For all datasets except snoRNA, we assembled SGs from DGs obtained using the values in Table 2. To assemble SGs from DGs, we filtered out reads having either arm longer than some *p*^th^ percentile arm length threshold and removed empty SGs, i.e. SGs for which all reads were eliminated. We then tested the stem formation potential of the RNA sequences spanned by the reads in each SG using ViennaRNA [20] at each percentile arm length threshold *p*, as described in the Methods section. We tested a range of percentile threshold *p*, ranging from 10^th^ to 90^th^ percentile at 10^th^ percentile increments. Dataset snoRNA was tested using both clustering methods over the full range of *t*_o_ and, where appropriate, ranges of *t*_eig_. For snoRNA, all clustering parameters and the full range of percentile thresholds were tested.

We refer to the resultant set of stem structures as CRSSANT SG base pairs, and compared them to the known base pairs. For each set of CRSSANT SG base pairs produced at different values of *p* (and, in the case of snoRNA, at all possible clustering parameter values), we recorded the following values: arm cutoff length, the total number of CRSSANT SG base pairs, number of base pairs that are shared between both the set of known base pairs and the set of CRSSANT SG base pairs, and the base pair recall (defined as the ratio of shared base pairs to known base pairs). The percentile threshold *p* for which the base pair recall is maximized was recorded as the optimal percentile cutoff, *p*_opt_. This rule is used to determine *p*_opt_, because while it is crucial to maximize the number of shared base pairs, it is also important to not penalize potentially novel base pairs, i.e. base pairs that are present in the set of CRSSANT SG base pairs but not in the known base pair set.

These results are summarized in Table 3. Complete SG base-pair results are in Additional file 2. The values in Table 3 recorded for snoRNA are those for which the number of shared base pairs was maximized over all clustering conditions and percentile thresholds. The CRSSANT SGs were found to generally have a larger number of base pairs than the known base pair set (Additional file 2, Additional file 4). However, the recall was found to cover a wide range, from a little under one-tenth for snRNA U4atac to nearly three-quarters for snRNA U5.

**Table 3.** Summary of results for the base pair comparison test. Optimal percentile cutoff *p*_opt_ and resultant arm cutoff length are reported for each dataset. The number of known base pairs, the number of base pairs in the CRSSANT SG stems and the number of base pairs shared between the known base pair and CRSSANT SG sets are also reported, alongside the base pair recall.

| dataset | gene | $p_{\text{opt}}$ | arm<br>cutoff<br>length | known<br>base pairs | CRSSANT<br>SG base<br>pairs | shared<br>base pairs | base pair<br>recall |
| --- | --- | --- | --- | --- | --- | --- | --- |
| RMRP | RMRP | 90 | 38 | 69 | 389 | 23 | 0.33 |
| RPPH1 | RPPH1 | 30 | 24 | 72 | 318 | 2 | 0.03 |
| rRNA | 18S | 40 | 28 | 477 | 1,058 | 110 | 0.23 |
|  | 5.8S | 20 | 24 | 20 | 66 | 7 | 0.35 |
|  | 28S | 40 | 26 | 1,495 | 1,812 | 340 | 0.23 |
| snRNA | U2 | 50 | 21 | 42 | 85 | 17 | 0.40 |
|  | U4 | 90 | 30 | 36 | 49 | 18 | 0.50 |
|  | U6 | 50 | 18 | 29 | 15 | 11 | 0.38 |
|  | U5 | 90 | 25 | 30 | 24 | 22 | 0.73 |
|  | U4atac | 90 | 27 | 21 | 26 | 2 | 0.10 |
|  | U6atac | 60 | 32 | 28 | 35 | 5 | 0.18 |
| snoRNA | U3 | 70 | 25 | 76 | 1,193 | 31 | 0.41 |

### Execution time

CRSSANT is fast and efficient at analyzing PARIS reads data. For all 12 datasets tested, the DG clustering step of the pipeline took less than four minutes to complete, and generating all nine SG output files took less than half a minute. Pipeline execution times under optimal DG clustering parameters are listed in Table 4.

**Table 4.** Execution times for running the CRSSANT pipeline on all datasets, separated into DG clustering and SG generation times.

| Dataset | Gene | DG clustering<br>[mm:ss] | SG generation<br>[ss] |
| --- | --- | --- | --- |
| RMRP | RMRP | 00:04 | 1 |
| RPPH1 | RPPH1 | 00:04 | 2 |
| rRNA | 18S | 00:54 | 19 |
|  | 5.8S | 00:01 | <1 |
|  | 28S | 00:11 | 19 |
| snRNA | U2 | 00:01 | <1 |
|  | U4 | 02:40 | <1 |
|  | U6 | 00:01 | <1 |
|  | U5 | 00:23 | <1 |
|  | U4atac | <00:01 | <1 |
|  | U6atac | <00:01 | <1 |
| snoRNA | U3 | 03:07 | 20 |

## Discussion

The results of the CRSSANT DG assignment validation tests set the default parameter values for both clustering methods, and instituted the use of spectral clustering as the default clustering method. There are two additional motivations for using spectral clustering as the default clustering method, rather than cliques-finding. First, the cliques-finding algorithm can be extremely time intensive, depending on the exact conditions of the dataset. We found that in certain instances, e.g. small nuclear ribonucleic acid spliceosmal RNA U4, the cliques-finding algorithm was prohibitively slow at certain overlap threshold and required nearly half an hour to complete (data not shown). In contrast, spectral clustering never required more than four minutes for any combination of clustering parameters. In addition to time constraints, the cliques-finding method tends to discard more reads than does the spectral clustering method. Ideally, the pipeline should not discard more reads than are necessary to avoid nonoverlapping reads in the same DG, which we found to be more often achieved using spectral clustering. Altogether, the variety of results shown in Table 2— notably, the number of nonzero non-overlapping reads for a number of the datasets—highlights the fact that although the CRSSANT method is able to automate otherwise laborious tasks, the user must still apply their own intuition to fine-tune the pipeline’s outputs. As a result, the pipeline is designed for customization. Users have the option of specifying different clustering parameter values. Furthermore, the pipeline checks for non-overlapping reads in all DGs. Whenever a DG containing non-overlapping reads is created, the pipeline automatically aborts, and provides general suggestions for improving clustering performance. These design decisions allow room for the user to interpret the results and experiment with different clustering settings.

User intuition is also required for interpreting the stem folding test results and output files. Despite the ease of predicting secondary structure from sequence alone, it is known that the accuracy of these algorithms is limited at best, especially with regard to predicting alternative structures [9]. In particular, though these algorithms are rooted in rigorous thermodynamic or comparative models, the computational complexity required to execute them often restricts the results to simplistic conformations. As a result, despite the relatively low number of shared base pairs between CRSSANT SG stems and known structures, the large number of novel base pairs in the CRSSANT SG sets demonstrates the potential for the PARIS assay to discover novel structures and interactions that have remained undetected by other means. This is bolstered by the low average distance between basic CRSSANT SG sets and known stem structures, described in Additional file 3 and shown in Additional file 4.

The wide variety of *p* selected as *p*_opt_ in the direct base pair tests further suggest that it is difficult to choose a good default percentile threshold a priori. This again implies that the best use of the CRSSANT pipeline may be to discover novel RNA stem formations by comparing to known base-pairing sets, rather than de-novo exploration of a completely unknown RNA structure. As a result, the CRSSANT pipeline outputs, by default, all nine sets of SG files: one SG base-pairing file, one SG arcs file and one SG auxiliary file for each percentile threshold ranging from 10 to 90, inclusive, in increments of 10, in order to allow CRSSANT users to apply their biological intuition to selecting the best SG stem set for their experimental purposes.

We acknowledge that the tests performed here are rather brute-force, but given the small volume of manually-curated DG assignment sets in existence and known structures found to overlap with CRSSANT structures, it is difficult to perform any kind of bootstrapping. We envision that default pipeline parameters can be updated as the PARIS method is adopted more widely, and note that the option to specify different pipeline parameters was built into CRSSANT to address this very issue.

Altogether, we have shown that the CRSSANT pipeline automates the otherwise laborious tasks of reads clustering and testing the structure-forming capabilities of gene regions implied by the clusters. However, it is still up to the user to fine-tune the desired output, and to interpret the biological significance of what the pipeline produces.

## Conclusions

We have presented the CRSSANT analysis pipeline which integrates network analysis techniques with state-of-the-art RNA structure prediction software in order to facilitate analysis of PARIS assay data towards the goal of secondary structure maps of RNA in the form of predicted base pair lists. We validated the read clustering and stem testing functionalities of CRSSANT against manually-labeled DG sets and known RNA structures over a range of human gene datasets including a variety of gene lengths and types. We also verified that CRSSANT’s computational requirements are reasonable.

We envision CRSSANT to be adopted as the default method of analyzing PARIS assay data, which is sure to itself be widely adopted as a method of in vivo elucidation of alternative structure and long-distance interaction formation in RNA molecules. CRSSANT is designed to not only automate clustering and structure prediction for the user, but also to streamline downstream analysis and experimentation. The files output by CRSSANT concisely summarize information that is crucial to the RNA structural biologist, and are prepared in file formats commonly used by the structural biology community in order to facilitate cross-platform analysis. Further improvements on CRSSANT include parameter tuning with the help of additional datasetes, adding the ability to analyze extremely large reads datasets with sampling, and implementing methods to further decrease runtime and improve operation efficiency.

## Methods

In this section, we outline the methods of the CRSSANT pipeline, which consist of the following components: data pre-processing, creating a network representation of an input PARIS reads set, network clustering, DG analysis, SG analysis and production of output files.

### Data pre-processing

The mapped reads accepted as input to the CRSSANT pipeline are assumed to be formatted in the SAM file format [19]. In order for the reads to be clustered, they must be first converted into 4-tuples of read arm start and stop indices. This is done using the read start position and its CIGAR alignment string, which encodes the transformations performed on the read during alignment. However, not all reads are readily converted due to variability in ligation. During the ligation step in the PARIS method, ligation may occur in two different ways. Ligation between the 3′ end of the left stem arm and the 5′ end of the right stem arms result in “normal” gapped reads. Ligation may occur at opposite ends of the stem, between the 5′ end of the left arm and the 3′ end of the right arm, resulting in “chiastic” reads. Due to the ability of RNA to form higher-level structures between different molecules in addition to intramolecular base-pairing interactions, chiastic reads may comprise sequences from two different strands or even genomic regions (i.e. chromosomes).

For simplicity, the CRSSANT pipeline operates on the assumption that all gapped reads in an input SAM file map to the same genomic region and strand. This assumption means that only chiastic reads from the same genomic region and strand are analyzed. Chiastic reads are output by the STAR aligner in a separate file and impossible to be represented with a single CIGAR string and read start position. Instead, every chiastic read is represented with two lines per read, each containing alignment information for the left and right read arms. The pre-processing module of the CRSSANT pipeline identifies chiastic reads that have both arms mapped to the same strand in the same genomic region, and integrates the two pieces of arm mapping information into a single SAM file line. These integrated lines are then appended to the SAM file containing all normal, non-chiastic reads. Finally, both normal and chiastic reads are converted into 4-tuples that can be used in a network representation.

### Network representation of PARIS reads

The reads are transformed into a network representation using the Python package NetworkX [12]. Each vertex in the network represents a single read. Consider a pair of reads *r*_1_, *r*_2_, each with left (*l*) and right (*r*) arm start and stop positions indexed by 0 and 1, respectively. Read *i* is represented by the 4-tuple of arm start and stop positions: (*a*_*i,l*,0_, *a*_*i,l*,1_, *a*_*i,r*,0_, *a*_*i,r*,1_).

To determine whether the network vertices representing the read pair *r*_1_, *r*_2_ are connected via an edge, left and right overlaps *o_j_*(*r*_1_, *r*_2_) and spans *s_j_*(*r*_1_, *r*_2_), *j* ∈ {*l, r*} of the read pair are calculated. Overlap and span are defined as:

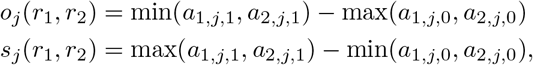

and are depicted in Figure 3.

**Fig. 3.**
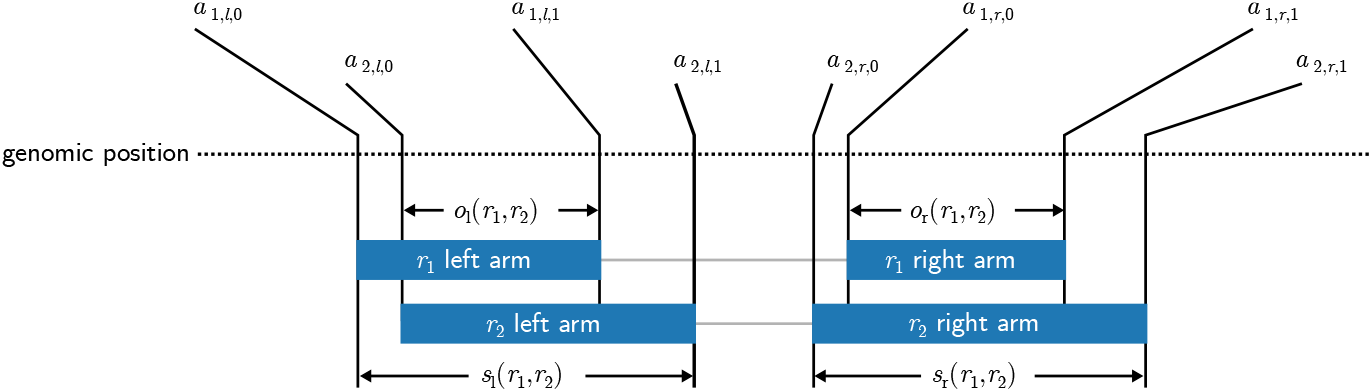
Overlap and span calculation for a pair of reads. Two reads *r*_1_ and *r*_2_ each comprising a left and right arm (solid blue bars), share left and right overlaps *o_l_*, *o_r_*, respectively and left and right spans *s_l_, s_r_*, respectively. The arm start and stop positions of read *i* are represented by the 4-tuple (*a*_*i,l*,0_, *a*_*i,l*,1_, *a*_*i,r*,0_, *a*_*i,r*,1_).

Note that the ratio of overlap to span is less than or equal to 0 if there is there is no overlap between arms, and is exactly 1 if the arms overlap completely and have the same length. An edge is drawn between the vertices representing reads *r*_1_, *r*_2_ in the network if both left and right overlap ratios *o_j_*(*r*_1_, *r*_2_)/*s_j_*(*r*_1_, *r*_2_) exceed some overlap threshold *t*_o_. This rule makes intuitive sense from a biological perspective: in order to have originated from the same stem structure, two reads must have at least some overlapping portion in both arms. The sum of the right and left overlap ratios of reads *r*_1_, *r*_2_ is recorded as the weight of the edge connecting the vertices representing reads.

To decrease graphing time, reads are ordered by all four arm start and stop positions. For a given read ordering “primary” read—one half of a potential read pair—is selected, and a “secondary” read—the second half of a potential read pair—is drawn sequentially from the remaining ordered reads. As long as the secondary read overlaps the primary read in both arms, the secondary read is added as a vertex to the network and the next read is selected as a candidate secondary read. However, once a candidate secondary read is found to share no overlap with the primary read, the remaining reads in the ordering are skipped, and reads from the next ordering are considered sequentially. This procedure of ordered traversals over all ordered reads avoids an exhaustive pairwise search over non-overlapping reads and increases pipeline efficiency.

We refer to the resulting network comprising the set of vertices *V* and the set of edges *E* as the weighted reads graph *G* = (*V, E*).

### Network clustering

The graph comprising all reads typically contains multiple subgraphs of connected components, which each represent groups of overlapping reads that do not overlap with other groups (i.e., that overlap less than overlap threshold *t*_o_). However, the subgraphs themselves may sometimes be further broken down into distinct clusters of reads that are more similar to each other than they are to other reads in the subgraph. To account for this possibility, the CRSSANT pipeline first partitions the weighted reads graph *G* into subgraphs of connected components, and then extracts clusters using two possible deterministic clustering methods: cliques-finding and spectral clustering.

Each method has its benefits and drawbacks. By definition, all vertices in a clique are fully connected, i.e. each vertex is accessible from every other vertex. In the biological setting, this can be interpreted as: all reads whose network representations comprise a clique must overlap in both the left and right arms beyond threshold *t*_o_. Intuitively, this implies that these reads are all highly similar, and that the read arm start and stop positions have low variability. Similarly, the arm start and stop positions of the DG comprising these reads can be obtained with high confidence. However, the requirement of full connectivity also has the undesirable potential to exclude a large number of reads which could discard valuable sequencing information. Furthermore, the problem of finding a maximal clique is often very computationally demanding [5].

Spectral clustering, on the other hand, does not require full connectivity between all vertices assigned to the same cluster, and thus provides a more flexible method of grouping vertices in a graph. In addition, it is simple to implement, can typically be solved efficiently using modern software, and often outperforms simple clustering algorithms like *k*-means clustering [32]. However, implementing spectral clustering depends on carefully choosing heuristics for a particular problem setting.

In the following subsections, we describe the two clustering methods and relevant parameters. The clustering parameter that is shared by both methods is the overlap threshold *t*_o_, which strongly affects subgraphs and, subsequently, the final DGs that PARIS reads are clustered into using cliques-finding or spectral clustering. In general, a large *t*_o_ results in a larger number of subgraphs containing fewer reads, since the reads must overlap more before an edge is drawn between their respective nodes. Exactly how the number of subgraphs and their composition affects the outcome of each clustering method is discussed in each of the following subsections.

#### Clustering method 1: Cliques

This method, an adaptation of [35], is built into the core NetworkX library and was used off-the-shelf. It divides each subgraph into cliques, or sets of nodes such that all nodes in the set share an edge with all other nodes. However, this implementation identifies all possible cliques that exist in the subgraph, meaning that a single read may exist in multiple cliques. This is in conflict with our goal of matching each read to a single DG. To deal with this ambiguity, we filter the list of all possible cliques using a greedy approach: cliques are sorted in descending order based on the number of reads in each clique, and candidate cliques are kept only if the set of kept cliques do not contain any of the reads in the candidate clique. The final set of cliques that are kept are declared to be the DGs. This method tends to discard a large number of reads, but has the benefit of resulting in DGs with highly similar reads, i.e. that share significant overlaps in both arms.

#### Clustering method 2: Spectral clustering

The spectral clustering method of cluster extraction is adapted from the Shi and Malik method described in [32], which we outline here briefly. For each subgraph *G_s_* = (*V_s_, E_s_*) containing *m* > 1 vertices, we calculate the *m* × *m* dimensional degree matrix *D* whose components *d_i,j_* are

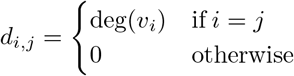

∀*v_i_* ∈ *G_s_*. The degree deg(*v_i_*) of a vertex is the number of times an edge terminates at that vertex. In terms of the PARIS reads, deg(*v_i_*) represents the number of reads overlapping the read represented by *v_i_* with overlap ratio exceeding *t*_o_. We also calculate the *m* × *m* dimensional unnormalized Laplacian matrix *L* of *G_s_* as

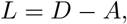

where *A* is the adjacency matrix of *G*_s_ with components *A_i,j_* = **1**_*v_i_,v_j_*_, *a_i,j_* ∈ {0, 1} indicating whether or not there exists an edge between vertices *v_i_* and *v_j_*. In terms of the PARIS reads, *A_i,j_* indicates whether or not the sum of the overlap ratios of both arms of *v_i_* and *v_j_* exceed *t*_o_. With *D* and *L*, we solve the generalized eigenproblem *Lv* = *λDv* using an eigengap heuristic based on the sorted eigenvalues and their corresponding eigenvectors to identify clustering parameter *k*, the number of clusters contained within the subgraph.

The eigengap heuristic chooses *k* such that the first *k* eigenvalues are “relatively small,” and the gap between the *k*^th^ eigenvalue *λ_k_* and the (*k* + 1)^th^ eigenvalue *λ*_*k*+1_ is “relatively large.” In practice, the variability of reads assigned to the same subgraph can result in eigenvalues that are not well separated, making it hard to choose the first relatively large eigengap. This choice is further complicated by the presence of large eigengaps among larger eigenvalues, which disqualifies decision rules based on a global analysis of all eigenvalues. To address these challenges, our implementation of the eigengap heuristic considers only the first *n*_eig_ eigenvalues. Then, the *k* – 1 eigengaps are calculated. For each eigengap we calculate what we call an “eigenratio,” i.e. the *i*^th^ eigenratio is the ratio between the *i*^th^ eigengap and the median of the preceding *i* – 1 eigengaps. The first eigenratio is simply the first eigengap. Finally, *k* is determined to be one less than the index of the first eigenratio that exceeds an eigenratio threshold *t*_eig_.

Intuitively, the goal of clustering is to create groups such that the edges connecting different groups have very low weights, while the edges connecting vertices within a group have high weight. Note that the Laplacian matrix *L* may be thought of as quantifying all edges emanating “out” from vertices, which are exactly the components that are used as the basis of clustering. Thus, solving the generalized eigenproblem involving the Laplacian *L* and identifying the first *k* largest eigengaps may be interpreted as the solution to the principal components problem of identifying the *k* dimensions which provide optimal separation of vertex groups.

In practice, we observed that PARIS data tended to result in first eigengaps (the difference between the first and second eigenvalues) that were small, and whose ratios rarely exceeded *t*_eig_. This resulted in a preponderance of *k* = 0 or *k* ≥ 3, even in situations where a human arbiter would have decided on *k* = 2. To allow for the possibility of splitting subgraphs into just two DGs, we added a second component to the eigengap heuristic that first checks if the magnitude of the second eigenvalue is large (empirically, greater than 1), and then if the remaining *n*_eig_ − 2 eigenvalues are much smaller than the second eigenvalue (empirically, if the median of the remaining eigenvalues is at least an order of magnitude smaller than the magnitude of the second eigenvalue). If these conditions are met, then *k* is set to 2. Once *k* is chosen, we use the first *k* eigenvectors to perform k-means clustering [15] on the subgraph vertices. Each resulting cluster is a DG.

Spectral clustering parameters *n*_eig_ and *t*_eig_ have opposite effects on DG clustering. A larger number of candidate eigenvalues, *n*_eig_, results in more eigengaps considered during execution of the eigengap heuristic. Considering more eigen-gaps increases the possibility of choosing a larger *k*, which results in splitting the subgraph into a larger number of DG clusters, each containing fewer reads.

On the other hand, increasing the eigengap threshold *t*_eig_ tends to result in smaller *k* which, in turn, results in splitting the subgraph into a smaller number of DG clusters. Each of these clusters will contain more reads than those that would have otherwise resulted from splitting the subgraph into a larger number of reads based on a smaller *t*_eig_.

### DG analysis

#### DG filtering

After reads are grouped into DGs, the DGs are filtered to remove low-quality DGs, and to arrive at a final set of DGs. Singleton DGs containing only a single read are eliminated and the read is excluded from further analysis. This filtering criterion creates a direct relationship between the graph and clustering parameters *t*_o_, *n*_eig_ and *t*_eig_ and the final number of DGs and final number of reads included for subsequent analysis. When the parameters are chosen such that there are fewer reads allocated to each DG, this increases the possibility that there exist singleton DGs, and in turn increases the possibility that more reads are eliminated from later analysis steps.

In addition to eliminating singleton DGs and reads, DGs are also checked for duplicate reads. If any DG contains only reads that are all identical in sequence, the reads are deemed to be duplicates. The DG is again eliminated, and the duplicate reads are excluded from further analysis.

#### DG attribute calculations

After DGs are filtered, various biologically significant attributes for each DG are calculated. The attributes are: number of reads, arm start and stop positions, coverage and non-overlapping group.

Since the PARIS method produces one read per RNA stem structure, the number of reads in each DG approximately corresponds to the cellular abundance of the stem structure represented by the DG. Potential DGs with very few reads may either represent structures with low abundance, or they may be clusters of reads that were incorrectly mapped due to sequencing errors. Thus, the number of reads making up a DG are recorded as an important attribute.

The 4-tuple of DG arm start and stop positions are calculated as the median of arm start and stop positions of all reads in the DG, and are an approximation of the gene sequence underlying the reads that engages in stem structure formation.

The coverage of DG *i* is defined in [24] to be

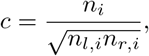

where *n_i_* is the total number of reads in DG *i, n_l,i_* is the number of reads whose left arms overlap the left arm of DG *i* and *n_r_i__* is the number of reads whose right arms overlap the right arm of DG *i*. Coverage is a score with range (0, 1], and is an approximation of how much the genomic region spanned by a DG engages in the formation of multiple RNA structures. A coverage score equal to the maximum value of 1 indicates that the genomic region spanned by the DG gives rise to only one RNA stem structure. On the other hand, a low coverage close to zero suggests that the genomic region spanned by the DG produces numerous RNA stem structures, in other words, that the genomic region engages in formation of multiple alternative RNA stem structures.

Finally, for each DG a non-overlapping group (NG) is calculated. DGs are assigned to NGs such that there are no overlaps between any of the reads in all DGs assigned to a given NG. NG assignments allow DGs to be easily viewed in genomic visualization software, like the popular Interactive Genomics Viewer [29].

### SG analysis

The ultimate goal of clustering PARIS reads is to identify regions of a gene that undergo base-pairing to form RNA stem structures. In order to identify these regions, DGs that pass the quality filter are further processed to produce stem groups (SGs) summarizing the characteristics of the stem structures.

#### SG assembly

The first step is to assemble SGs by filtering from the DGs all reads that are above a certain length threshold. The filtering process as follows: all reads from DGs that passed the quality filter are aggregated, and the *p*^th^ percentile arm length is calculated. Then, any read comprising an arm that is longer than the *p*^th^ percentile arm length is discarded.

The SGs are then reviewed to remove any SGs that may have had all reads eliminated during the reads filtering process.

#### SG attribute calculations

As with DGs, the number of reads in each SG and SG arm start and stop positions are reported. Since SGs approximate regions of genes participating in RNA stem structure formation, we additionally use state-of-the-art RNA folding software to test the stem formation potential of the regions spanned by SGs. For each SG tested, we report the base pairs composing the stem, the number of potential psoralen cross-linking sites, and the length of the stem.

As with DGs, the number of reads in an SG are recorded. Since a number of reads may have been removed from the corresponding DG during SG assembly, it is important to re-calculate the number of reads in all SGs.

Unlike the arm start and stop positions of DGs, those for SGs are calculated by considering the minimum and maximum arm start and stop positions, respectively. Biologically, reads grouped together in the same SG can be thought of as randomly digested sections of the same stem structure formed by different RNA molecules. In other words, each read reveals some valuable information about the underlying stem structure. As a result, we take the minimum left arm start position of all reads to be the left arm start position of the SG. Similarly, we take the maximum right arm stop position of all reads to be the right arm stop position of the SG. However, the left arm stop and right arm start positions are not so simple, since in order for the SG arms to hybridize, they must not overlap. To account for this fact, plus the possibility of overhanging single-stranded regions between the arms, we calculate the SG’s left arm stop position to be the minimum of the maximum of the left arm stop positions and the minimum of the right arm start positions. Similarly, the SG’s right arm start position is taken to be the maximum of the maximum of the left arm stop positions and the minimum of the right arm stop positions.

The potential of the RNA sequences spanned by the arms of the reads making up each SG to form a stable RNA stfem structure is tested using the ViennaRNA software package’s RNAfold method, as outlined in the Methods section. RNAfold contains a number of RNA folding test functions, and we use the `fold_compound` function because it both allows the setting of hard constraints on RNA base-pairing consistent with stem formation, and also returns biologically significant information about the stem. To ensure that ViennaRNA produces canonical Watson-Crick base-paired stems from the SG’s left and right arm sequences without allowing portions of each arm to basepair with themselves, we set the hard constraints that a base at genomic position *i* in the SG’s left arm must pair with a base at position *j*, where *i* < *j*, and vice versa for the bases in the SG’s right arm. Stems with a negative minimum free energy are considered to be molecularly stable; all other stems are discarded. For the recorded stems, we save the returned base pairs in the stem (reported using dot-bracket notation [14]), the estimated minimum free energy of the folded stem and the number of paired bases in the folded stem as attributes of the SG.

Finally, we also examine the SG arm sequences for the presence of staggered uridine bases and staggered uridine-cytosine base pairs, which are the preferred substrates for psoralen cross-linking [7]. The number of urdine-uridine and uridine-cytosine base pairs is also recorded as an SG attribute.

### Output files

The CRSSANT pipeline analyzes reads by gene pairs and produces up to five types of output files for reads overlapping a given gene pair (*g*_1_, *g*_2_). Gene pairs may be either automatically generated from the set of all genes represented by the reads in the input SAM file, or may be specified individually by the user. Restricting analysis to a single gene is represented by the gene pair (*g*_1_, *g*_1_). For each gene pair, only reads from the input SAM file whose left arms map to *g*_1_ and whose right arms map to *g*_2_ are analyzed. If reads from the gene pair form valid DGs that do not contain any non-overlapping reads, the two output files are produced: a SAM file and a BED file [1]. It is important to check for nonoverlapping reads within the same DG, since the presence of a read left arm begins downstream of other reads’ right arms in the same DG is invalid—there is no biological basis for such reads being clustered together and no evidence that they originated in the same genomic region. As a result, if any DGs contain non-overlapping reads, DG results are not written to output files.

All reads successfully clustered into DGs that passed the filtering step are written to the output SAM file. This SAM file is identical to the input SAM file, with two additional annotations for DGs and NGs. For each read assigned to DG *N*, the string `DG : i : N` is appended to the read line in the SAM file. The string `NG : i : M` is also appended to the read line, where *M* is the NG to which DG *N* belongs. Attributes of these DGs are written to the output BED file which includes the genomic region of the DG, the DG identification number, coverage, number of reads, arm start and stop positions and arm lengths.

Next, SG results are checked. If SGs are successfully assembled from the DGs, i.e., if not all SGs are filtered out after the percentile thresholding step, an additional three output files are produced: an SG base-pairing file, an SG arcs file and an auxiliary file containing miscellaneous SG information. The SG base-pairing and SG arcs files are formatted as BED files, and are intended to help the user visualize the RNA structures implied by SGs that were successfully assembled from DGs. The SG base-pairing file specifies arcs for each base pair in the stems formed by the SGs. The SG arc file specifies arcs between the centers of each SG’s left and right arms. Finally, the SG auxiliary file contains all remaining SG attributes that were calulated for each SG, including SG ID number and coverage of the corresponding SG, number of reads, number of uridine-uridine and uridine-cytosine base pairs and number of base pairs in the stem. The SG auxiliary file also contains the minimum, maximum and standard deviation of each arm’s start and stop positions, a quick summary of the reads composing each SG.

