## Additional file 3 for "Cross-linked RNA Secondary Structure Analysis using Network Techniques"

### Basic stems analysis for “Cross-linked RNA Secondary Structure Analysis using Network Techniques”

When testing CRSSANT stem groups (SGs), we found that ViennaRNA folding results are sensitive to comparatively small changes in arm cutoff length. This is because changes in arm cutoff length determine whether entire reads are excluded or not, which can significantly change the arm start and stop positions of assembled SGs and, in turn, affect the possible base pairs that form in the CRSSANT SG stems.

To account for these variations, and also for the fact that the ViennaRNA structure prediction software relies upon necessarily simplified models of molecular folding, we performed an additional evaluation on the CRSSANT SG stems. In the “SG structure validation and prediction” subsection of the results, we compared individual base pairs between structure sets and the SG stem structures. Here, we compare entire regions where base-pairing occurs. This type of analysis allows us to move beyond a granular comparison on the level of individual base pairs towards a higher-level understanding of which areas in a gene, with some spatial flexibility, are engaged in forming structures and long-range interactions.

To do this, we grouped base pairs into collections of consecutive base pairs along each strand. This aggregation method easily identifies classic RNA stems comprising Watson-Crick base pairs, but does not account for more complex RNA structures such as stems with bulges, stems with internal loops, and pseudoknots. As a result, we refer to these collections of base pairs as “basic” stems.

For each basic stem formed from CRSSANT SG base pair files and ground truth base pair files, the start and stop indices were recorded and used to calculate a number of quantities for each percentile threshold. First, the number of SG basic stems sharing a nonzero overlap in both arms with at least one ground truth basic stem was tallied. This number provides a general idea of how many of the total number of SG basic stems were validated by basic stems in the ground truth sets, and also how many SG basic stems were previously unknown. The number of ground truth stems that share a nonzero overlap in both arms with at least one basic CRSSANT SG stem is also recorded, since stem overlaps do not have a 1:1 correspondence as base pairs do. Next, the average stem distance $\bar{d}$ is recorded. The stem distance $d(s_1, s_2)$ is defined to be the Euclidean distance between any basic CRSSANT SG stem $s_1$ and any basic ground truth stem $s_2$ with which $s_1$ shares a nonzero overlap in both arms. Stem distance between stems $s_1$ and $s_2$ with arm start and stop indices $(a_{i,l,0}, a_{i,l,1}, a_{i,r,0}, a_{i,r,1})$ with $i \in \{1, 2\}$, and $l, r$ denoting left and right arm indices, respectively, and $0, 1$ denoting arm start and stop indices, respectively, is calculated as:

$$d(s_1, s_2) = \sqrt{(a_{1,l,0} - a_{2,l,0})^2 + (a_{1,l,1} - a_{2,l,1})^2 + (a_{1,r,0} - a_{2,r,0})^2 + (a_{1,r,1} - a_{2,r,1})^2}.$$

The stem distance has units in bases and may thus be easily interpreted.

The results from the stem comparison tests are summarized in Additional file 4. As was the case for the direct base pair evaluations, all data sets except for snoRNA were clustered using the clustering methods and parameters specified in Table 2 in the main text. As with the direct base pair comparison test, the number of overlapping stems is relatively small compared to the number of SG and ground truth stems. However, the average distance between overlapping stems ranges between 0—a perfect match—to less than six bases, a very small difference in stems.
